# Evidence that inconsistent gene prediction can mislead analysis of algal genomes

**DOI:** 10.1101/690040

**Authors:** Yibi Chen, Raúl A. González-Pech, Timothy G. Stephens, Debashish Bhattacharya, Cheong Xin Chan

**Affiliations:** Institute for Molecular Bioscience, University of Queensland, Brisbane, QLD 4072, Australia; School of Chemistry and Molecular Biosciences, University of Queensland, Brisbane, QLD 4072, Australia; Department of Biochemistry and Microbiology, Rutgers University, New Brunswick, NJ 08901, USA

## Abstract

Comparative algal genomics often relies on predicted gene models from *de novo* assembled genomes. However, the artifacts introduced by different gene-prediction approaches, and their impact on comparative genomic analysis, remain poorly understood. Here, using available genome data from six dinoflagellate species in Symbiodiniaceae, we identified potential methodological biases in the published gene models that were predicted using different approaches. We developed and applied a comprehensive customized workflow to predict genes from these genomes. The observed variation among predicted gene models resulting from our workflow agreed with current understanding of phylogenetic relationships among these taxa, whereas those published earlier were largely biased by the distinct approaches used in each instance. Importantly, these biases mislead the inference of homologous gene families and synteny among genomes, thus impacting biological interpretation of these data. Our results demonstrate that a consistent gene-prediction approach is critical for comparative genomics, particularly for non-model algal genomes.

---

We implemented a customized, comprehensive workflow to predict protein-coding genes in six published draft Symbiodiniaceae genomes: *Breviolum minutum* (Shoguchi et al. 2013), *Symbiodinium tridacnidorum, Cladocopium* C92 (Shoguchi et al. 2018), *Symbiodinium microadriaticum* (Aranda et al. 2016), *Cladocopium goreaui* and *Fugacium kawagutii* (Liu et al. 2018). These draft genomes, generated largely using short-read sequence data, remain fragmented (e.g. N50 lengths range from 98.0 Kb for *C. goreaui* to 573.5 Kb for *S. microadriaticum*); we treated these genome assemblies independently as is standard practice. The published gene models from these four studies were predicted using three different approaches: (a) *ab initio* using AUGUSTUS (Stanke et al. 2006) guided by transcriptome data (Shoguchi et al. 2013, Shoguchi et al. 2018), (b) *ab initio* using AUGUSTUS guided by a more-stringent selection of genes (Aranda et al. 2016), and (c) a more-thorough approach incorporating evidence from transcriptomes, machine learning tools, homology to known sequences and *ab initio* methods (Liu et al. 2018). Because repetitive regions are commonly removed prior to gene prediction, multi-copy genes are sometimes mis-identified as repeats and excluded from the final gene models. To address this issue, we adapted the workflow from Liu et al. (2018) to ignore inferred repeats in the final step that integrates multiple evidence sources using EVidenceModeler (Haas et al. 2008). To minimize potential contaminants in the published draft genomes and their impact on gene prediction, we identified and removed genome scaffolds that share high similarity (BLASTn, *E* ≤ 10-20, bit-score ≥ 1000, query cover ≥ 5%) to bacterial, archaeal and viral genome sequences in the RefSeq database (release 88), adopting a similar approach to Liu et al. (2018). We then compared, for each genome, the published gene models in the remaining scaffolds against the predicted gene models in these same scaffolds using our approach. Specifically, we assessed metrics of gene models, and the inference of homologous gene families and conserved synteny within a phylogenetic context.

For simplicity, hereinafter we refer to the published gene models as *α* genes, and those predicted in this study as *β* genes. Compared to *α* genes, the structure of *β* genes (based on the distribution of intron lengths) resembles more closely the structure of dinoflagellate genes inferred using transcriptome data (Figure S1). These results suggest that *β* genes are likely more biologically realistic. Variation between *α* and *β* genes was assessed using ten metrics: number of predicted genes per genome, average gene length, number of exons per genome, average exon length, number of introns per genome, average intron length, proportion of splice-donor site motifs (GT, GC or GA), number of intergenic regions, and average length of intergenic regions.

As shown in Table S1, the metrics for *α* and *β* genes differed substantially. The number of *α* genes per genome was much higher in some cases and showed greater variation (mean 48,050; standard deviation 16,741) than that of *β* genes (mean 32,819; standard deviation 7567). This is likely due to the more-stringent criteria used by our workflow to delineate protein-coding genes. The larger variation in the number of *α* genes is likely due to biases arising from the distinct prediction methods and not assembly artifacts, because the same genome assembly for each species was used to independently derive *α* and *β* genes. Most predicted genes (>60% genes in each genome) were supported by transcriptome evidence (BLASTn, *E* ≤ 10-10). In some cases, *β* genes have stronger transcriptome support than *α* genes; e.g. 82.6% compared to 66.9% in *S. tridacnidorum*, and 78.4% compared to 61.9% in *Cladocopium* C92 (Table S1).

Variation in the ten observed metrics among *α* and *β* genes was also assessed using PCA (Fig. 1a). The *α* genes are more widespread along principal component 1 (PC1, between −0.54 and 0.46), with those based on AUGUSTUS-predominant workflows distinctly separated (PC1 < −0.19; Fig. 1a). The *β* genes are distributed more narrowly on PC1 (between 0 and 0.27) and more widely along principal component 2 (PC2; between −0.55 and 0.20). Interestingly, the distribution of genes along PC2 exhibits a pattern that is consistent with our current understanding of the phylogeny of these six species (Fig. 1b). Specifically, the *Symbiodinium* species are clearly separated from the others along PC2 (Fig. 1a) and the two *Cladocopium* species are clustered more closely based on *β*, rather than *α* genes. Therefore, PC1 (explaining 51.46% of the variance) largely reflects the variation introduced by distinct gene prediction methods, whereas the distribution along PC2 (explaining 25.91% of the variance) is likely attributable to the phylogeny of these species. This result suggests that variation among *α* genes is predominantly due to methodological biases, and that these biases are larger compared to those of *β* genes. Variation in the latter appears to be more biologically relevant and consistent with Symbiodiniaceae evolution.

**Fig. 1.**
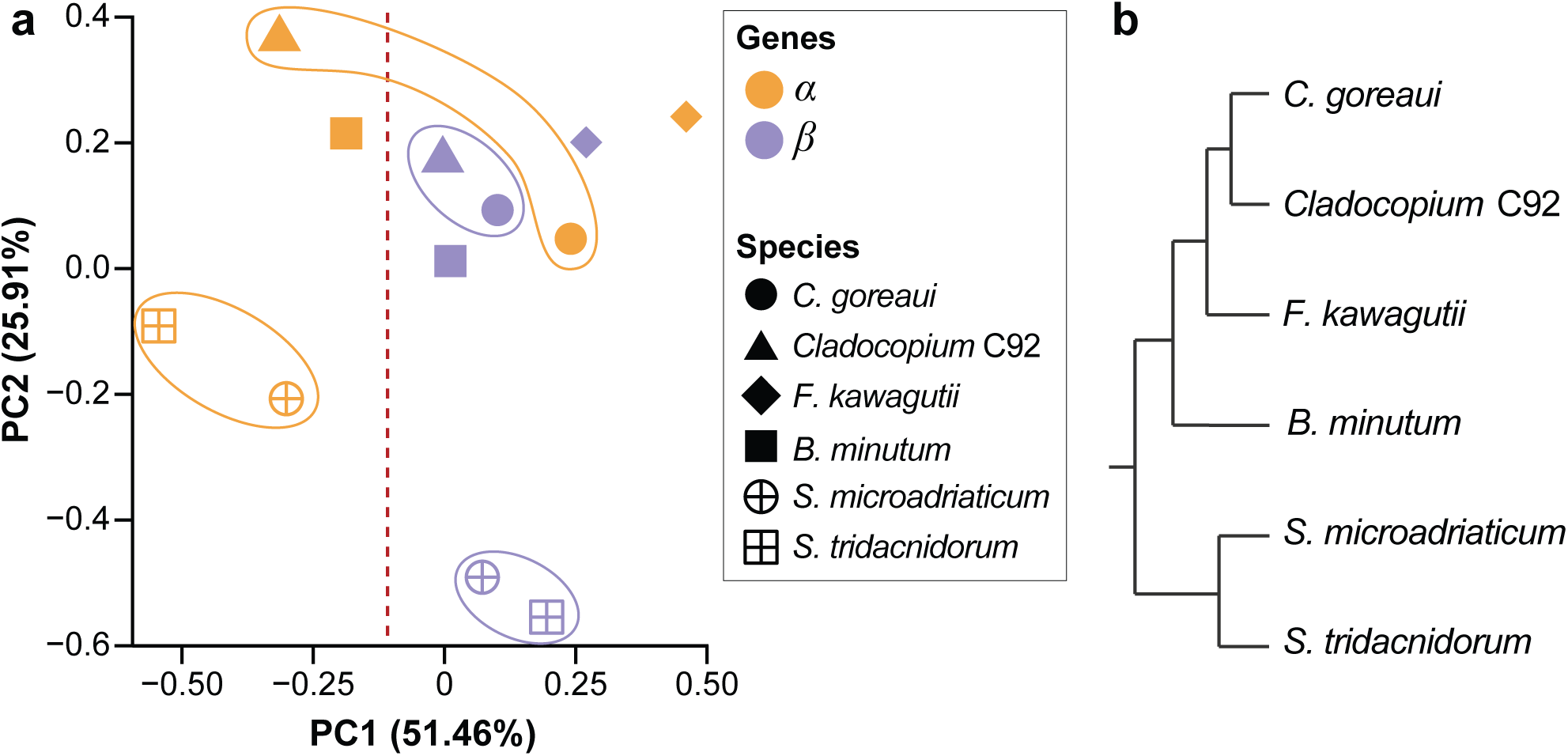
Gene metrics of *α* and *β* genes from six Symbiodiniaceae genomes. (a) Principal Component Analysis plot based on ten metrics of the predicted gene models, shown for the *α* genes in orange, and the *β* genes in purple, for each of the six genomes (noted in different symbols) as indicated in the legend. The two *Cladocopium* species and the two *Symbiodinium* species were highlighted for clarity. (b) A tree topology depicting the phylogenetic relationship among the six taxa, based on (LaJeunesse et al. 2018).

Genomes that are phylogenetically closely related are expected to share greater synteny than those that are more distantly related. Here, we defined a collinear syntenic gene block as a region common to two genomes in which five or more genes are coded in the same order and orientation. These gene blocks were identified using SynChro (Drillon et al. 2014) at *Delta* = 4. Overall, 421 collinear syntenic blocks (implicating 2454 genes) between any genome-pairs were identified among *α* genes, compared to 450 blocks (implicating 2728 genes) among *β* genes (Figs. 2a and 2b). Based on the *α* genes comparison (Fig. 2a), *S. microadriaticum* and *S. tridacnidorum* shared the largest number of syntenic blocks (130; 760 genes), whereas *S. microadriaticum* and *F. kawagutii* shared the fewest (1; 6 genes). Surprisingly, *S. tridacnidorum* and *Cladocopium* C92 shared 38 blocks (222 genes). This close relationship is not evident between any other pair of genomes from these two genera (e.g. only 3 blocks implicating 15 genes between *S. microadriaticum* and *C. goreaui*), and is even closer than the relationship between the two *Cladocopium* species (i.e. *C. goreaui* and C92: 33 blocks, 187 genes). In an independent analysis, the unexpectedly high conserved synteny between *S. tridacnidorum* and *Cladocopium* C92 was attributed to inflated evidence support from isoforms of similar *α* genes (as predicted by AUGUSTUS), and the structural configuration (i.e. combination of exons) among *α* genes that is distinct from that among *β* genes. This observation may be explained by the fact that *α* genes from these two genomes were predicted using the same method (Shoguchi et al. 2018). In contrast, based on the *β* genes comparison (Fig. 2b), the number of syntenic blocks shared between any *Symbiodinium* and *Cladocopium* species did not vary to the same extent; e.g. 7 blocks (38 genes) between *S. tridacnidorum* and *Cladocopium* C92, and 10 blocks (55 genes) between *S. microadriaticum* and *C. goreaui*. The number of *β* genes implicated in blocks shared by these two genera is also smaller than those between the two *Cladocopium* species (263 genes in 48 blocks), consistent with their closer phylogenetic relationship.

**Fig. 2.**
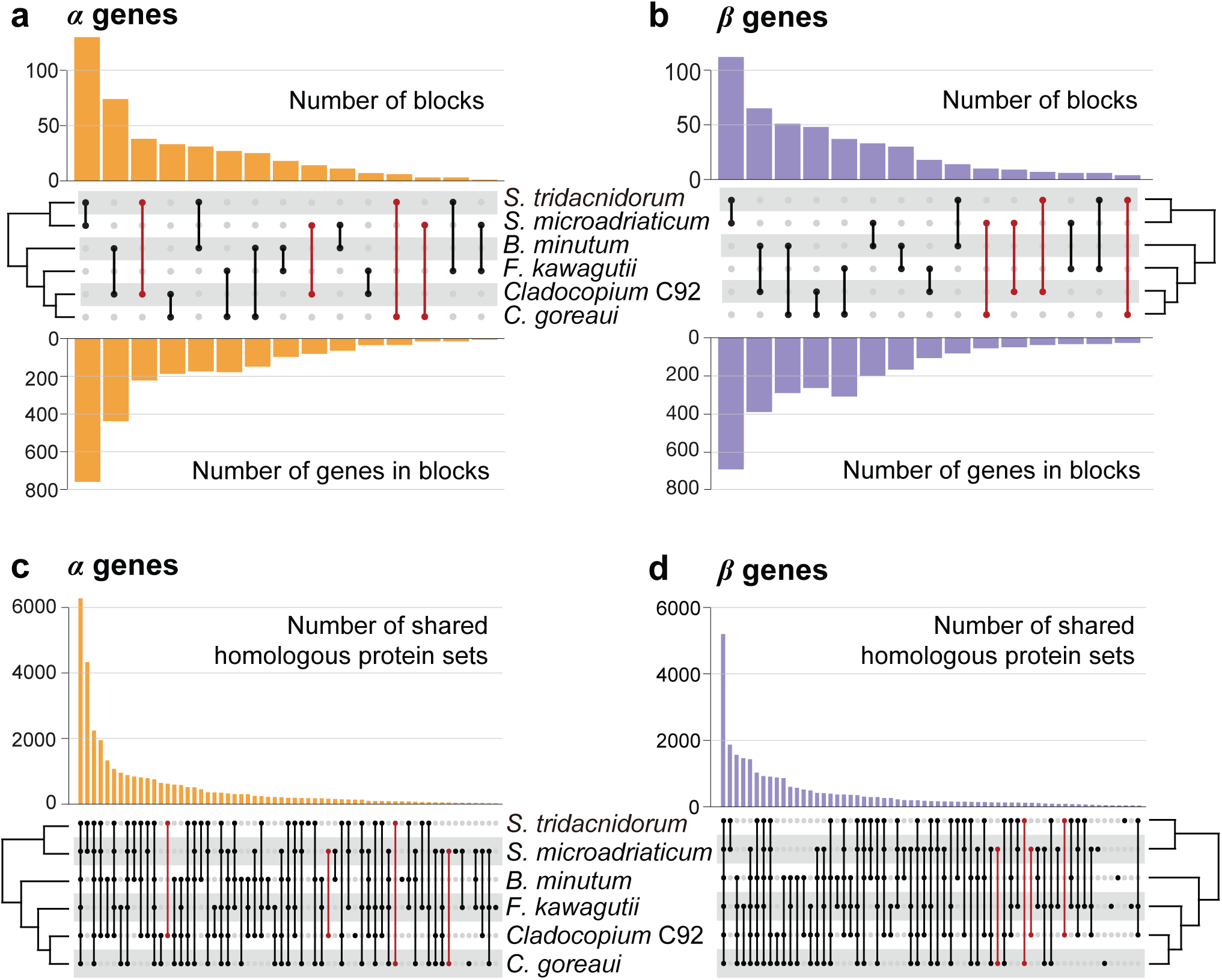
Conserved synteny and homologous sets among six Symbiodiniaceae genomes. The number of collinear syntenic gene blocks between each genome-pair is shown for those inferred based on (a) *α* and (b) *β* genes; the upper bar chart shows the number of blocks, the lower bar chart shows the number of implicated genes in these blocks, and the middle panel shows the genome-pairs corresponding to each bar with a line joining the dots that represent the implicated taxa. The number of homologous sets inferred from (c) *α* and (d) *β* genes is shown, in which the taxa represented in the set corresponding to each bar are indicated in the bottom panel. The most remarkable differences between (a) and (b), and (c) and (d), focusing on *Symbiodinium* and *Cladocopium* species, are highlighted in red.

To assess the impact of methodological biases on the delineation of homologous gene families, Orthofinder v2.3.1 (Emms & Kelly 2018) was used to infer “orthogroups” from protein sequences (i.e. homologous protein sets) encoded by the *α* and *β* genes (Figs 2c and 2d). More homologous sets were inferred among the *α* genes (33,580) than among the *β* genes (26,924), likely due to the higher number of *α* genes in all genomes. Genomes from closely related taxa are expected to share more homologous sequences (and therefore more sets) than those that are phylogenetically distant. Most of the identified homologous sets (6431 from *α* genes, 5217 from *β* genes) contained sequences from all analyzed taxa; these represent core gene families of Symbiodiniaceae. Similar to the results of the synteny analysis described above, the pattern of homologous sets shared between members from *Symbiodinium* and *Cladocopium* varies among the *α* genes (Fig. 2c). For instance, 638 homologous sets are shared only between *S. tridacnidorum* and *Cladocopium* C92, compared to 89 between *C. goreaui* and *S. tridacnidorum*. In contrast, the corresponding number of homologous sets inferred based on *β* genes are closer to each other (Fig. 2d); i.e. 92 between *S. tridacnidorum* and *Cladocopium* C92, and 123 between *C. goreaui* and *S. tridacnidorum*.

Our results indicate that comparative genomics using the *α* genes (i.e. simply based on published gene models) could lead to the inference that *S. tridacnidorum* and *Cladocopium* C92 are more closely related with each other than is each of them with other isolates in their corresponding genus. The bias introduced by different gene-prediction approaches can significantly impact downstream comparative genomic analyses and lead to incorrect biological interpretations. We therefore urge the research community to consider a consistent gene-prediction workflow when pursuing comparative genomics, particularly among highly divergent, non-model algal genomes. Although we only considered dinoflagellate genomes from a single family in this study, the implication of our results can be applied more broadly to all other non-model eukaryote genomes.

## Acknowledgements

TGS is supported by an Australian Government Research Training Program Scholarship. RAGP is supported by an International Postgraduate Research Scholarship and a University of Queensland Centenary Scholarship. TGS was supported by an Australian Government Research Training Program Scholarship. This work was supported by two Australian Research Council grants (DP150101875 awarded to Mark Ragan, CXC and DB, and DP190102474 awarded to CXC and DB), and the computational resources of the Australian National Computational Infrastructure (NCI) Facility through the NCI Merit Allocation Scheme (project d85) awarded to CXC.

## Data accessibility

All genome data (after removal of microbial contaminants), and all predicted gene models from this study are available at: https://cloudstor.aarnet.edu.au/plus/s/JXALPndBKLNYgF9

## Author contribution

YC, RAGP and CXC conceived the study and designed the experiments. YC conducted all computational analyses. All authors analyzed and interpreted the results. YC and RAGP prepared all figures, tables, and the first draft of this manuscript. YC, TGS and RAGP provided analytical tools and scripts. All authors wrote, reviewed, commented on and approved the final manuscript.

## Supplementary Information

**Fig. S1. Distribution of intron lengths in predicted genes from six Symbiodiniaceae genomes.** In each graph, the distribution of intron lengths among α genes (orange line), among *β* genes (purple line), and among transcript-based gene models (predicted using PASA v2.3.3 and TransDecoder v5.2.0; red dashed line) are shown. The transcript-based gene models were considered as a proxy for true gene structure.

**Table S1. Metrics of predicted gene models in genomes of Symbiodiniaceae.**

