## Supplementary Figure S1 and Table S1 for "Evidence that inconsistent gene prediction can mislead analysis of algal genomes"

**Supplementary Information**

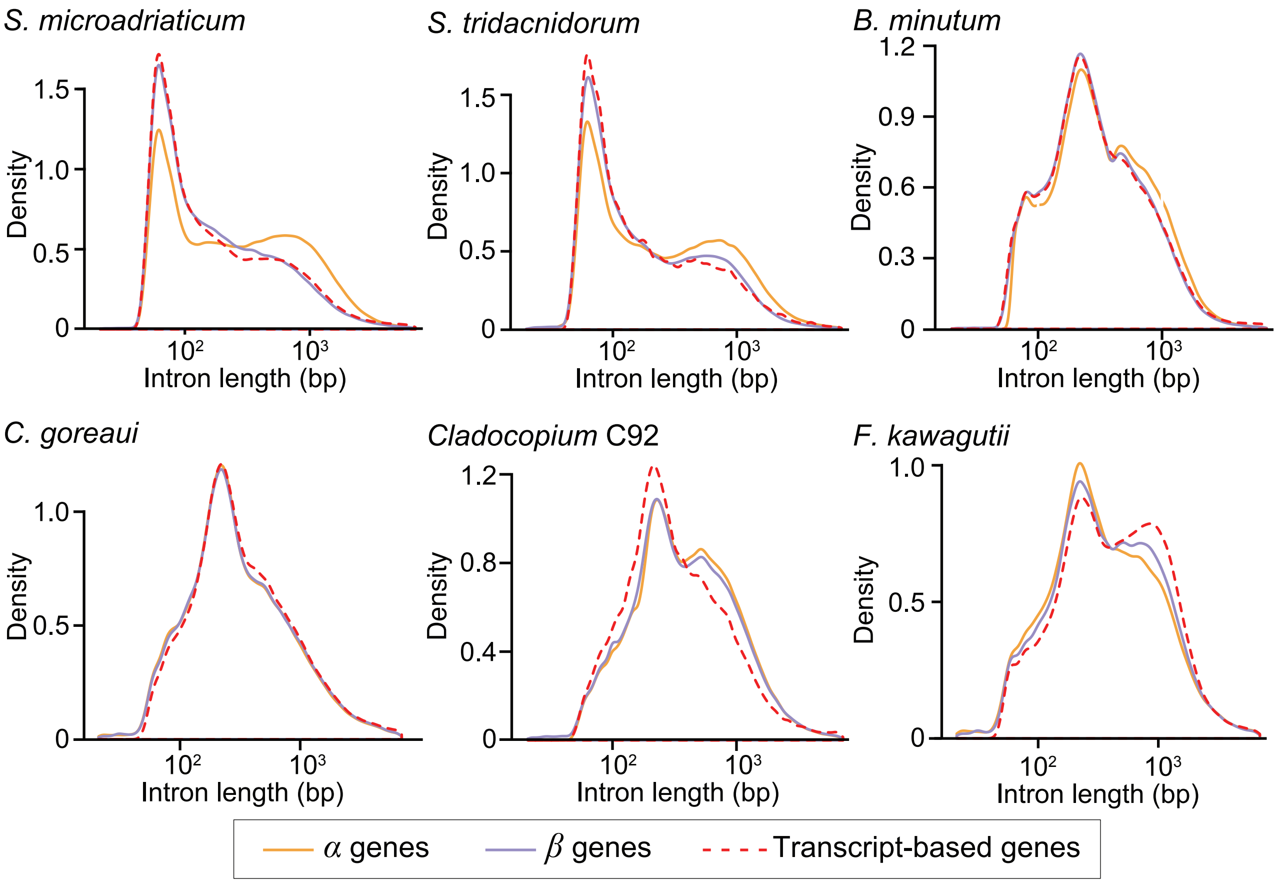


**Fig. S1. Distribution of intron lengths in predicted genes from six Symbiodiniaceae genomes.** In each graph, the distribution of intron lengths among *α* genes (orange line), among *β* genes (purple line), and among transcript-based gene models (predicted using PASA v2.3.3 and TransDecoder v5.2.0; red dashed line) are shown. The transcript-based gene models were considered as a proxy for true gene structure.

**Table S1. Metrics of predicted gene models in genomes of Symbiodiniaceae.**

| Metric | | *B. minutum* | | *Cladocopium* C92 | | *S. microadriaticum* | | *S. tridacnidorum* | | *C. goreaui* | | *F. kawagutii* | |
|---|---|---|---|---|---|---|---|---|---|---|---|---|---|
| | | *α* | *β* | *α* | *β* | *α* | *β* | *α* | *β* | *α* | *β* | *α* | *β* |
| **Number of genes** | | 41,920 | 32,803 | 65,832 | 33,421 | 49,012 | 29,728 | 69,018 | 25,808 | 35,913 | 47,114 | 26,609 | 33,341 |
| **Average gene length (bp)** | | 11,962 | 10,069 | 8193 | 13,207 | 12,918 | 9281 | 8835 | 6467 | 6967 | 7595 | 6507 | 8615 |
| **Average CDS length (bp)** | | 2318 | 1930 | 1870 | 2201 | 2392 | 2218 | 2093 | 1425 | 1766 | 1902 | 1736 | 1932 |
| **Transcript support (%)** | | 91.5 | 89.4 | 61.9 | 78.4 | 76.5 | 79.2 | 66.9 | 82.6 | 66.8 | 63.6 | 63.9 | 65.2 |
| **Genes with introns (%)** | | 95.4 | 93.7 | 80.4 | 97.6 | 98.2 | 95.7 | 83.1 | 88.2 | 92.9 | 94.6 | 94.0 | 95.5 |
| **Number of exons per gene** | | 23.5 | 19.0 | 16.0 | 18.6 | 21.9 | 19.2 | 22.7 | 12.9 | 10.0 | 10.9 | 8.7 | 11.3 |
| **Average exon length (bp)** | | 98.6 | 101.4 | 117.0 | 118.1 | 109.3 | 115.4 | 92.0 | 110.3 | 175.9 | 174.0 | 199.5 | 171.5 |
| **Number of introns per gene** | | 22.4 | 18.0 | 14.8 | 17.6 | 20.9 | 18.2 | 21.5 | 11.9 | 9.0 | 9.9 | 7.7 | 10.3 |
| **Average intron length (bp)** | | 516.8 | 451.3 | 607.6 | 623.8 | 504.6 | 387.9 | 509.7 | 423.1 | 575.1 | 573.3 | 619.4 | 651.4 |
| **Splice donor motif (%)** | **GT** | 48.2 | 43.9 | 47.0 | 41.6 | 26.0 | 26.0 | 33.8 | 24.6 | 36.3 | 44.3 | 39.5 | 40.5 |
| | **GC** | 36.6 | 37.9 | 41.4 | 41.4 | 52.1 | 54.1 | 55.5 | 57.2 | 43.0 | 36.3 | 39.8 | 38.3 |
| | **GA** | 15.2 | 18.2 | 11.7 | 17.0 | 21.8 | 19.9 | 10.7 | 18.2 | 20.6 | 19.4 | 20.7 | 21.1 |
| **Splice acceptor with AGG motif (%)** | | 95.6 | 96.4 | 95.1 | 95.2 | 96.2 | 96.4 | 95.4 | 97.0 | 95.5 | 95.6 | 94.4 | 95.0 |
| **Number of intergenic regions** | | 32,429 | 25,383 | 58,946 | 28,504 | 42,232 | 26,713 | 55,407 | 18,677 | 23,898 | 32,295 | 21,709 | 28,061 |
| **Average intergenic region length (bp)** | | 1993 | 5984 | 2062 | 6337 | 3627 | 15,110 | 1808 | 13,545 | 10,627 | 8076 | 23,042 | 17,141 |
